# Improved genome sequence and annotation of *Cutaneotrichosporon oleaginosus* ATCC 20509

**DOI:** 10.1101/2024.03.20.585711

**Authors:** Bart Nijsse, Zeynep Efsun Duman-Özdamar, Janine A. C. Verbokkem, Derek Butler, Maria Suarez-Diez, Mattijs K. Julsing

## Abstract

*Cutaneotrichosporon oleaginosus* is an oleaginous yeast with a high content of fatty acids and can accumulate more than 40% of its weight in lipids. It can grow on a wide range of carbon sources and side streams such as crude glycerol. The genome sequence of *C. oleaginosus* ATCC 20509 is reported here to contribute to its development as a biotechnological platform for producing microbial oils.

## Introduction

*Cutaneotrichosporon oleaginosus* ATCC 20509 is an oleaginous yeast that can accumulate more than 40% lipids (w/w) of its biomass under nutrient-limiting conditions [1]. *C. oleaginosus* has also been known as *Apiotrichum curvatum, Cryptococcus curvatus, Trichosporon cutaneum, Trichosporon oleaginosus*, and *Cutaneotrichosporon curvatum* and it is reported among the top five most studied oil-producing yeasts with around 20 % of the publications [2]. Under nitrogen-limiting conditions, the produced fatty acid composition has been reported to be 30 % palmitic acid (C16:0), 8 % stearic acid (C18:0), 47 % oleic acid (C18:1), and 12 % linoleic acid (C18:2) which is comparable to that of palm oil [3]. Therefore the microbial oil produced by *C. oleaginosus* has strong potential as a sustainable alternative in various industrial applications such as biofuels, feed, chemicals, personal care, and cosmetic products. Furthermore, these yeasts can use a broad range of carbon sources, such as glucose, xylose, glycerol, sucrose, and lactose[4, 5]. They can also use more complex and inexpensive side streams such as crude glycerol from bio-ethanol production or whey permeate as a feedstock [6, 1]. Due to the mentioned advantages, *C. oleaginosus* has been flagged as a promising alternative to develop an industrial production process for microbial oil. The genome of this organism had already been sequenced [7]. Here, we present an updated sequence obtained combining short and long reads. RNA sequencing datasets have been incorporated into genome structural annotation to improve gene prediction.

## Methods

### Cultivation and DNA isolation C. oleaginosus

ATCC 20509 was grown on YPD broth (10 g/L yeast extract, 20 g/L peptone, 20 g/L glucose) at 30°C and 250 rpm for 20 hours. Cells were harvested and resuspended with 1:10 volumes of CutSmart Buffer (NEB, # B7204) and 5 µl of 100 mg/ml RNase-A solution (pre-heated at 100°C for 20 minutes). After 1 hour incubation at 37 °C cells were harvested and supernatant was mixed with 1:10 volume of 2M NaAC pH 5.0. 0.54 volume of isopropanol was added and it was placed at -20°C for 15 minutes. The sample was centrifuged at 13000 g for 15 minutes and washed twice with 1 ml 70% ethanol. Following the centrifugation for 5 minutes at 13000 g, the pellet was dried and dissolved in sterile Milli-Q.

### DNA sequencing and quality filtering

Single-end and paired-end sequence short reads were generated using the Illumina NovaSeq 6000 or MiSeq system, respectively The sequences generated with the MiSeq system were performed under accreditation according to the scope of BaseClear B.V. (L457; NEN-EN-ISO/IEC 17025). FASTQ read sequence files were generated using bcl2fastq2 version 2.18. The initial quality assessment was based on data passing the Illumina Chastity filtering. Subsequently, reads containing PhiX control signal were removed using an in-house filtering protocol. The quality of Illumina reads was improved by trimming off low-quality bases using BBDuk, which is a part of the BBMap suite version 36.77 [**?** ]. Long reads were sequenced using the PacBio Sequel system. The data collected from the PacBio Sequel instrument was processed and filtered using the SMRT Link software suite. Subreads shorter than 50 bp were discarded.

### Genome assembly

In the first step a draft genome was assembled. ABySS version 2.0.2 was used to assemble the high-quality reads were assembled into contigs [8]. Then, BLASR version 1.3.1 was used to map the long reads to the draft assembly [9]. These alignments were used to link the contigs and place them into scaffolds. Orientation, order, and distance between the contigs were estimated using SSPACE-LongRead version 1.0 [10]. Illumina reads were used to (partly) closed gapped regions within scaffolds using GapFiller version 1.10 [11]. Finally, possible assembly errors and nucleotide disagreements between the Illumina reads and scaffold sequences were corrected using Pilon version 1.21 [12]. The final assembly has an average coverage of 50.6 with the PacBio reads and 331 with the short Illumina reads.

### Genome annotation

RNA sequencing reads were retrieved from ENA study PRJEB73355 were quality-filtered and trimmed using fastp v0.20.0 [13]. Ribosomal RNA was removed using bbduk from the BBMap v38.57 using the available ribokmers.fa.gz file in the BBMap suite [14]. Read inspection was performed using FastQC v0.11.9 and MultiQC v1.8 [15, 16]. Filtered reads from all conditions were mapped to the genome with STAR v2.7.3a [17]. The resulting BAM file was used for gene prediction with FUNGAP v1.1.0 using the tools GeneMark-ES Suite v4.32, RepeatModeler-open v1.0.11 and BRAKER v1.9 [18, 19, 20, 21]. The BUSCO dataset was set to dikarya_odb9, the augustus_species parameter to “cryptococcus_neoformans_gattii” and the sister proteomes of two *Cutaneotrichosporon oleaginosum* species (GCF_001027345.1, GCA_008065305.1) and *Vanrija humicola* (GCA_008065275.1) were used as input. The resulted gff3 file was converted to an EMBL for ENA submission with EMBLmyGFF3 [22].

## Results

Illumina sequencing resulted in 22916792 short reads with an average quality (Q-score) of 35.86. In addition 528795 long reads were generated with a mean read length of 5122 bp and a maximum length of 86097 bp. The *de novo* hybrid assembly resulted in an assembled genome with 9 scaffolds (see Table 1) and 8132 protein coding genes. The genome sequence is submitted to ENA under project number PRJEB73837, assembly accession GCA_963992775 and scaffold accessions CAXDVN010000001-CAXDVN010000009.

**Table 1.**
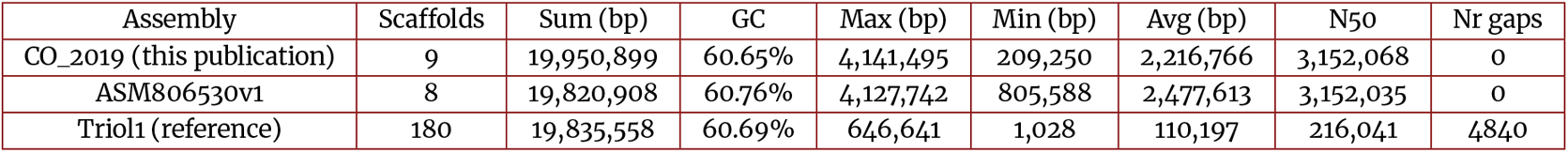
Genome assembly characteristics.

**Table 2.**
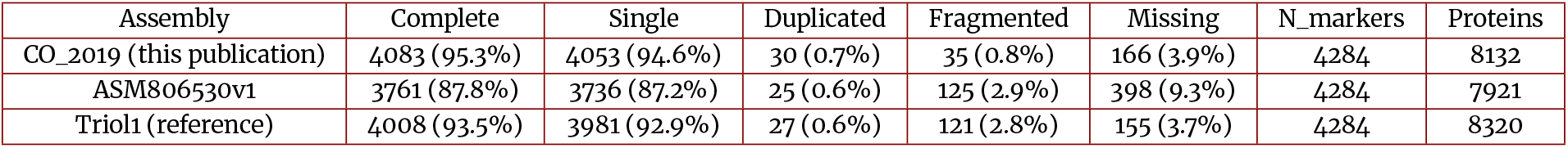
BUSCO scores with dataset: tremellomycetes_odb10 2024-01-08.

## References

1. Bracharz F, Beukhout T, Mehlmer N, Brück T. Opportunities and challenges in the development of Cutaneotrichosporon oleaginosus ATCC 20509 as a new cell factory for custom tailored microbial oils. Microb Cell Fact 2017;16.

2. Abeln F, Chuck CJ. The history, state of the art and future prospects for oleaginous yeast research. Microb Cell Fact 2021;20.

3. Duman-Özdamar ZE, Martins dos Santos VAP, Hugenholtz J, Suarez-Diez M. Tailoring and optimizing fatty acid production by oleaginous yeasts through the systematic exploration of their physiological fitness. Microb Cell Fact 2022;21.

4. Pham N, Reijnders M, Suarez-Diez NB M, Springer J, Eggink G, Schaap PJ. Genome-scale metabolic modeling underscores the potential of Cutaneotrichosporon oleaginosus ATCC 20509 as a cell factory for biofuel production. Biotechnol Biofuels 2021;14.

5. Shaigani P, Awad D, Redai V, Fuchs M, Haack M Mehlmer, et al. Oleaginous yeasts-substrate preference and lipid productivity: a view on the performance of microbial lipid producers. Microb Cell Fact 2021;20.

6. Cui Y, Blackburn JW, Liang Y. Fermentation optimization for the production of lipid by Cryptococcus curvatus: Use of response surface methodology. Biomass and Bioenergy 2012;47:410–417. https://www.sciencedirect.com/science/article/pii/S0961953412003571.

7. Close D, Ojumu J. Draft Genome Sequence of the Oleaginous Yeast <i>Cryptococcus curvatus</i> ATCC 20509. Genome Announcements 2016;4(6):10.1128/genomea.01235-16. https://journals.asm.org/doi/abs/10.1128/genomea.01235-16.

8. Jackman SD, Vandervalk BP, Mohamadi H, Chu J, Yeo S, Hammond SA, et al. ABySS 2.0: resource-efficient assembly of large genomes using a Bloom filter. Genome Research 2017 May;27(5):768–777. https://genome.cshlp.org/content/27/5/768, company: Cold Spring Harbor Laboratory Press Distributor: Cold Spring Harbor Laboratory Press Institution: Cold Spring Harbor Laboratory Press Label: Cold Spring Harbor Laboratory Press Publisher: Cold Spring Harbor Lab.

9. Chaisson MJ, Tesler G. Mapping single molecule sequencing reads using basic local alignment with successive refinement (BLASR): application and theory. BMC Bioinformatics 2012 Sep;13(1):238. 10.1186/1471-2105-13-238.

10. Boetzer M, Pirovano W. SSPACE-LongRead: scaffolding bacterial draft genomes using long read sequence information. BMC Bioinformatics 2014 Jun;15(1):211. 10.1186/1471-2105-15-211.

11. Boetzer M, Pirovano W. Toward almost closed genomes with GapFiller. Genome Biology 2012;13:R56. 10.1186/gb-2012-13-6-r56.

12. Walker BJ, Abeel T, Shea T, Priest M, Abouelliel A, Sakthikumar S, et al. Pilon: An Integrated Tool for Comprehensive Microbial Variant Detection and Genome Assembly Improvement. PLOS ONE 2014 Nov;9(11):e112963. https://journals.plos.org/plosone/article?id=10.1371/journal.pone.0112963, publisher: Public Library of Science.

13. Chen S, Zhou Y, Chen Y, Gu J. fastp: an ultra-fast all-in-one FASTQ preprocessor. Bioinformatics (Oxford, England) 2018 Sep;34(17):i884–i890.

14. Bushnell B, BBMap; 2014. https://sourceforge.net/projects/bbmap/.

15. Andrews S, FastQC: A Quality Control Tool for High Throughput Sequence Data; 2010. http://www.bioinformatics.babraham.ac.uk/projects/fastqc/.

16. Ewels P, Magnusson M, Lundin S, KÃ¤ller M. MultiQC: summarize analysis results for multiple tools and samples in a single report. Bioinformatics 2016 Oct;32(19):3047–3048. 10.1093/bioinformatics/btw354.

17. Dobin A, Davis CA, Schlesinger F, Drenkow J, Zaleski C, Jha S, et al. STAR: ultrafast universal RNA-seq aligner. Bioinformatics 2013 Jan;29(1):15–21. https://www.ncbi.nlm.nih.gov/pmc/articles/PMC3530905/.

18. Min B, Grigoriev IV, Choi IG. FunGAP: Fungal Genome Annotation Pipeline using evidence-based gene model evaluation. Bioinformatics 2017 Sep;33(18):2936–2937. 10.1093/bioinformatics/btx353.

19. Ter-Hovhannisyan V, Lomsadze A, Chernoff YO, Borodovsky M. Gene prediction in novel fungal genomes using an ab initio algorithm with unsupervised training. Genome Research 2008 Dec;18(12):1979–1990. https://genome.cshlp.org/content/18/12/1979, company: Cold Spring Harbor Laboratory Press Distributor: Cold Spring Harbor Laboratory Press Institution: Cold Spring Harbor Laboratory Press Label: Cold Spring Harbor Laboratory Press Publisher: Cold Spring Harbor Lab.

20. Flynn JM, Hubley R, Goubert C, Rosen J, Clark AG, Feschotte C, et al. RepeatModeler2 for automated genomic discovery of transposable element families. Proceedings of the National Academy of Sciences 2020 Apr;117(17):9451–9457. https://www.pnas.org/doi/full/10.1073/pnas.1921046117, publisher: Proceedings of the National Academy of Sciences.

21. Hoff KJ, Lomsadze A, Borodovsky M, Stanke M. Whole-Genome Annotation with BRAKER. Methods in Molecular Biology (Clifton, NJ) 2019;1962:65–95.

22. Norling M, Jareborg N, Dainat J. EMBLmyGFF3: a converter facilitating genome annotation submission to European Nucleotide Archive. BMC Research Notes 2018;11:1–5.

